# Structure and mechanism of a cyclic trinucleotide-activated bacterial endonuclease mediating bacteriophage immunity

**DOI:** 10.1101/694703

**Authors:** Rebecca K. Lau, Qiaozhen Ye, Lucas Patel, Kyle R. Berg, Ian T. Mathews, Jeramie D. Watrous, Aaron T. Whiteley, Brianna Lowey, John J. Mekalanos, Philip J. Kranzusch, Mohit Jain, Kevin D. Corbett

## Abstract

Bacteria possess an array of defenses against foreign invaders, including diverse nucleases that target and destroy the genome of an invading bacteriophage or foreign DNA element. A recently-described bacteriophage immunity pathway employs a cGAS/DncV-like nucleotidyltransferase to produce a cyclic tri-AMP second messenger, which activates the DNA endonuclease effector NucC. Here, we show that NucC is related to restriction enzymes but uniquely assembles into a homotrimer in solution. cAAA binding in a conserved allosteric pocket promotes assembly of two NucC trimers into a homohexamer competent for double-strand DNA cleavage. We propose that NucC mediates bacteriophage immunity either through global activation, causing altruistic cell death and abortive infection, or through local activation and targeted phage genome destruction. Finally, we identify NucC homologs in type III CRISPR-Cas systems, where they likely function as accessory nucleases activated by cyclic oligoadenylate second messengers synthesized by these systems’ effector complexes.

## Introduction

Bacteria are locked in a continual struggle to protect themselves and their genomes from foreign mobile genetic elements and bacteriophages, and as such have evolved sophisticated defenses against these invaders. Two well-characterized bacterial defense systems are restriction-modification systems and CRISPR-Cas systems, both of which rely on targeted nuclease activity to destroy foreign nucleic acids. In a typical restriction-modification system, a bacterium encodes a sequence-specific DNA endonuclease, and a corresponding DNA methylase that modifies the host genome to protect it against endonuclease cleavage (Arber and Linn, 1969; Wilson and Murray, 1991). While restriction-modification systems mount a standard response to all improperly modified DNA and as such can be likened to an “innate” immune system, the more recently-characterized CRISPR-Cas systems are more akin to an “adaptive” immune system. In these systems, effector nucleases are loaded with guide sequences that specifically target foreign nucleic acids (Koonin et al., 2017; Sorek et al., 2013; Wright et al., 2016). In type III CRISPR-Cas systems, the multi-protein effector complex not only recognizes and cleave foreign RNA and DNA (Koonin et al., 2017; Pyenson and Marraffini, 2017), but also synthesizes 3’-5’ linked cyclic oligoadenylate second messengers upon target binding (Kazlauskiene et al., 2017; Niewoehner et al., 2017). These second messengers in turn activate accessory RNases encoded in the same operons, including Csm6 (type IIIA systems) and Csx1 (type IIIB) (Han et al., 2018; Kazlauskiene et al., 2017; Niewoehner et al., 2017).

We recently characterized a newly-described bacterial defense pathway encoding a cGAS/DncV-like nucleotidyltransferase (CD-NTase) plus regulator proteins related to eukaryotic HORMA domain proteins and the AAA+ ATPase Pch2/TRIP13 (Ye et al., 2019). Our findings support a model in which detection of specific foreign peptides by bacterial HORMA domain proteins stimulates synthesis of a nucleotide second messenger, which activates one of several classes of effector proteins (Burroughs et al., 2015; Whiteley et al., 2019; Ye et al., 2019). We characterized two operons from patient-derived strains of *E. coli* and *Pseudomonas aeruginosa* whose CD-NTases synthesize the cyclic trinucleotide second messenger cyclic AMP-AMP-AMP (cAAA). Both operons also encode a restriction endonuclease-related effector protein termed NucC (<u>Nuc</u>lease, <u>C</u>D-NTase associated), which is activated by cAAA and required for bacteriophage immunity (Ye et al., 2019). How a restriction endonuclease-related enzyme can be controlled by second messenger binding, however, remains an important unanswered question.

Here, we show that NucC is a homotrimeric DNA endonuclease that binds the second messenger cAAA in a three-fold symmetric allosteric pocket. We show that nuclease activation involves the assembly of two NucC trimers into a homohexamer, which juxtaposes pairs of endonuclease active sites and promotes coordinated double-strand DNA cleavage. NucC is largely sequence non-specific and is not inhibited by DNA methylation, suggesting that high-level activation in cells may cause genome destruction and cell death. Finally, we identify type III CRISPR operons that encode NucC homologs in place of the accessory RNases Csm6/Csx1, defining a new family of type III CRISPR systems encoding a cyclic oligoadenylate-activated accessory DNase.

## Results

### Bacterial CD-NTase associated nucleases adopt a homotrimeric architecture

We have recently shown that a bacterial CD-NTase-encoding operon from a patient-derived strain of *E. coli* (MS115-1) provides immunity against bacteriophage infection, likely acting by sensing one or more phage proteins and synthesizing cAAA in response (Ye et al., 2019). The putative effector protein in this operon, which we term NucC (<u>Nuc</u>lease, <u>C</u>D-NTase-associated), is a cAAA-activated DNA endonuclease necessary for phage immunity (Ye et al., 2019). To understand the structure of NucC and its mechanism of activation, we overexpressed and purified *E. coli* MS115-1 (*Ec*) NucC, and a second NucC from a related operon found in *Pseudomonas aeruginosa* strain ATCC27853 (*Pa*) (Fig. 1A). Both *Ec* NucC and *Pa* NucC self-associate in yeast two-hybrid assays (Ye et al., 2019), and we used size exclusion chromatography coupled to multi-angle light scattering (SEC-MALS) to show that both proteins form homotrimers in solution (Fig. S1A-B). While several families of DNA exonucleases – including bacteriophage λ nuclease (Kovall and Matthews, 1997; Zhang et al., 2011) and the archaeal nuclease PhoExoI (Miyazono et al., 2015) – are known to form homotrimers, endonucleases are more commonly homodimeric, with two active sites that cleave opposite DNA strands to effect double-strand cleavage (Pingoud et al., 2005).

**Figure 1.**
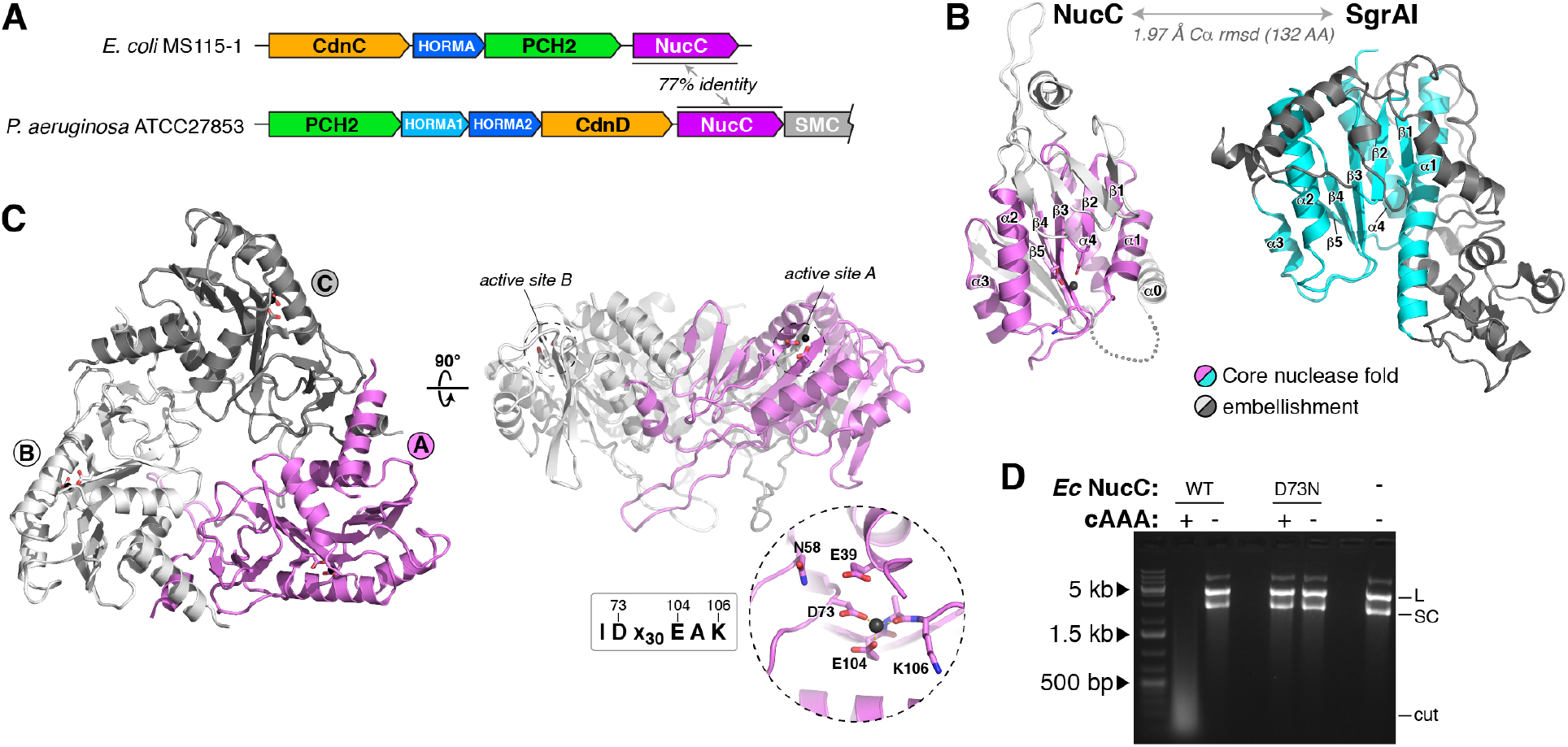
NucC is a homotrimeric relative of restriction endonucleases. **(A)** Schematic of CD-NTase operons from *E. coli* MS115-1 and *P. aeruginosa* ATCC27853 harboring NucC (Ye et al., 2019). **(B)** Structure of an *Ec* NucC monomer and the related restriction endonuclease SgrAI (PDB ID 3DW9; (Dunten et al., 2008)). Shared secondary structural elements in the core nuclease fold are colored violet (NucC) or cyan (SgrAI), and family-specific embellishments are shown in gray. **(C)** Structure of an *Ec* NucC trimer with monomers colored violet (monomer A), white (B), and gray (C). The side view (*right*) shows the active sites of the A and B monomers. *Inset:* closeup view of the NucC active site, with ordered Mg^2+^ ion shown as a black sphere. NucC possesses a conserved IDx_30_EAK active-site consensus motif (See Fig. S1C for comparison with the active site of SgrAI, and Fig. S1E for NucC sequence alignment). **(D)** Plasmid digestion assay with wild-type and active-site mutant D73N *Ec* NucC (10 nM), in the presence of cAAA (500 nM). “L” denotes linear plasmid, “SC” denotes closed-circular supercoiled plasmid, and “cut” denotes fully-digested DNA.

We crystallized and determined the structure of *Ec* NucC to a resolution of 1.75 Å (Table S1). In agreement with prior sequence analysis (Burroughs et al., 2015), the NucC monomer adopts a restriction endonuclease fold (Fig. 1B). A database search using the DALI structure-comparison server (Holm and Rosenström, 2010) revealed that NucC is structurally related to type II restriction endonucleases, including NgoMIV (Deibert et al., 2000), Kpn2I (PDB ID 6EKR; unpublished), and SgrAI (Dunten et al., 2008). Indeed, comparing NucC with a structure of SgrAI bound to DNA (Dunten et al., 2008) reveals an overall r.m.s.d. of 1.97 Å over 132 aligned Cα atoms (Fig. 1B and S1C-D). NucC also possesses a conserved active site motif of IDx_30_EAK (residues 72-106 in *Ec* NucC), with the conserved acidic residues Asp73 and Glu104 coordinating a Mg^2+^ ion in our structure (Fig. 1C). Mutation of Asp73 to Asn disrupts the endonuclease activity of NucC, demonstrating that DNA cleavage likely occurs through an equivalent mechanism as type II restriction endonucleases (Fig. 1D). We previously showed that the same mutation disrupts the ability of the *E. coli* MS115-1 CD-NTase operon to confer immunity against bacteriophage infection (Ye et al., 2019). Consistent with our SEC-MALS data, the structure of NucC reveals a tightly-packed homotrimer with an overall triangular architecture (Fig. 1C). The three active sites are arrayed on the outer edge of the trimer. The trimer interface of NucC is composed largely of elements that are not part of the core nuclease fold, but are rather embellishments not shared with dimeric restriction endonucleases (Fig. S1E).

### NucC binds cAAA in an allosteric pocket

We next tested binding of NucC to a range of dinucleotide and trinucleotide (Fig. S2) second messengers using isothermal titration calorimetry (ITC). We found that *Ec* NucC binds strongly to 3’,3’,3’cAAA (*Kd* = 0.7 μM), binds more weakly to both 3’,3’cyclic di-AMP (*Kd* = 2.6 μM) and linear di-AMP (5’-pApA; 4.4 μM), and does not bind 2’,3’cyclic di-AMP or AMP (Fig. S3). No second messenger bound with a stoichiometry higher than ~0.3 molecules per NucC monomer in our ITC assays, suggesting that the NucC trimer likely possesses a single second messenger binding site.

We next co-crystallized *Ec* NucC with 3’,3’,3’cAAA and determined the structure of the complex to a resolution of 1.66 Å (Table S1). The structure reveals that a single cAAA molecule binds in a conserved, three-fold symmetric allosteric pocket on the “bottom” of the NucC trimer (Fig. 2A). Each adenine base is recognized through a hydrogen bond between its C6 amine group and the main-chain carbonyl of Val227, and through π-stacking with Tyr81 (Fig. 2B-C). In addition, NucC residues Arg53 and Thr226 form a hydrogen-bond network with each phosphate group of the cyclic trinucleotide (Fig. 2B). Binding of cAAA induces a dramatic conformational change in an extended hairpin loop in NucC, which we term the “gate loop”, resulting in the closure of all three loops over the conserved pocket to completely enclose the bound cAAA molecule (Fig. 2D). Two aromatic residues, His136 and Tyr141, interact through π-stacking to stabilize the gate loop’s structure in both bound and unbound states. In the cAAA-bound structure, these two residues serve to “latch” the gate loops through tight association at the three-fold axis of the NucC trimer. Mutation of gate loop residues His136 or Tyr141, or cAAA-binding residues Arg53, Tyr81, or Thr226 completely disrupts NucC’s cAAA-activated endonuclease activity (Fig. 2F).

**Figure 2.**
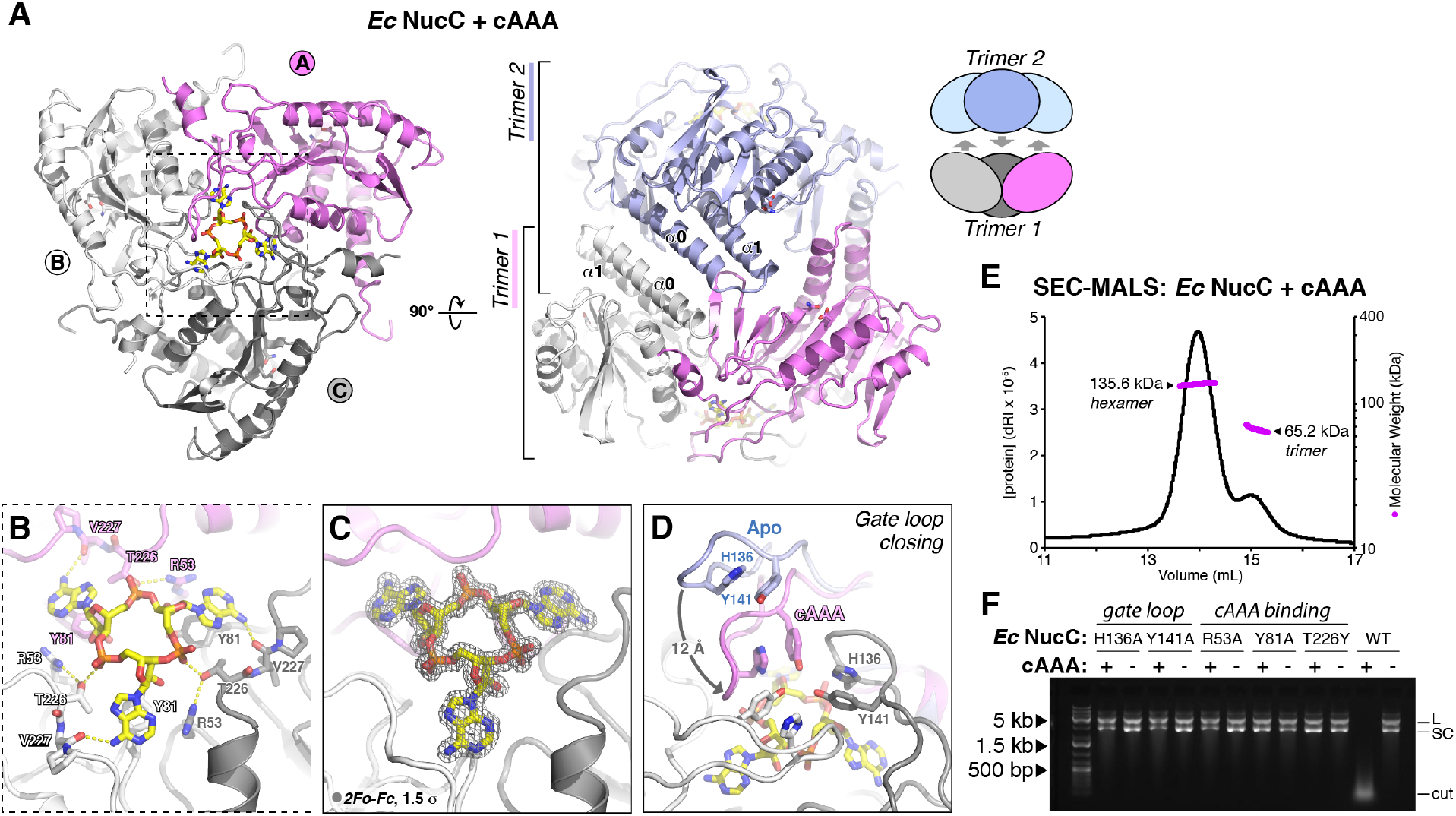
NucC binds cAAA in an allosteric pocket. **(A)** *Left:* Bottom view of a NucC trimer bound to cAAA, with subunits colored violet/gray/white as in Fig. 1C. *Right:* Side view of a NucC hexamer, assembled from two trimers of cAAA-bound NucC (one violet/gray/white, the second blue) interacting through packing of their α0 helices. See Fig. S4 for structure of cAAA-bound *Pa* NucC forming an equivalent hexamer. **(B)** Closeup view of cAAA binding NucC. **(C)** *2Fo-Fc* electron density for cAAA at 1.66 Å resolution, contoured at 1.5 σ. **(D)** Closeup view of gate loop closure upon cAAA binding. For one monomer, the gate loop is shown in both open (Apo, blue) and closed (cAAA-bound, violet) conformations, showing the ~12 Å motion between the two states. Pi stacking of “latch” residues H136 and Y141 (labeled in gray monomer) stabilizes gate loop conformation in both states. **(E)** Size exclusion chromatography coupled to multi-angle light scattering (SEC-MALS) for *Ec* NucC after pre-incubation with cAAA. While untreated *Ec* NucC shows a single peak corresponding to a homotrimer (Fig. S1A), cAAA treatment results in the appearance of a second peak corresponding to a homohexamer (calculated MW 160.3 kDa). **(F)** Plasmid digestion assay with wild-type and structure-based mutants of *Ec* NucC (10 nM) in the presence and absence of cAAA (100 nM).

### cAAA-mediated NucC hexamerization activates double-strand DNA cleavage

In the crystals of cAAA-bound *Ec* NucC, two NucC trimers associate to form a homohexamer through association of their α0 helices (Fig. 2A). This binding causes residues 23-34 linking α0 and α1, which are disordered in our NucC Apo structure and border the NucC active site, to become well-ordered in the cAAA-bound structure. We used SEC-MALS to confirm that both *Ec* NucC and *Pa* NucC form hexamers in solution when incubated with cAAA (Fig. 2E and Fig. S4). These data suggest that cAAA activates NucC’s nuclease activity in part by stimulating the formation of a NucC homohexamer.

We next modelled DNA binding by overlaying the NucC monomer with a structure of DNA-bound SgrAI (Dunten et al., 2008). Hexamer formation juxtaposes pairs of NucC active sites from opposite trimers ~25 Å apart (Fig. 3A), and we could model a continuous bent DNA duplex that contacts a pair of juxtaposed NucC active sites (Fig. 3B). In this model, the putative scissile phosphate groups are positioned two base pairs apart on opposite DNA strands (Fig. 3B), such that coordinated cleavage at these two sites would yield a double-strand break with a two-base 3’overhang.

**Figure 3.**
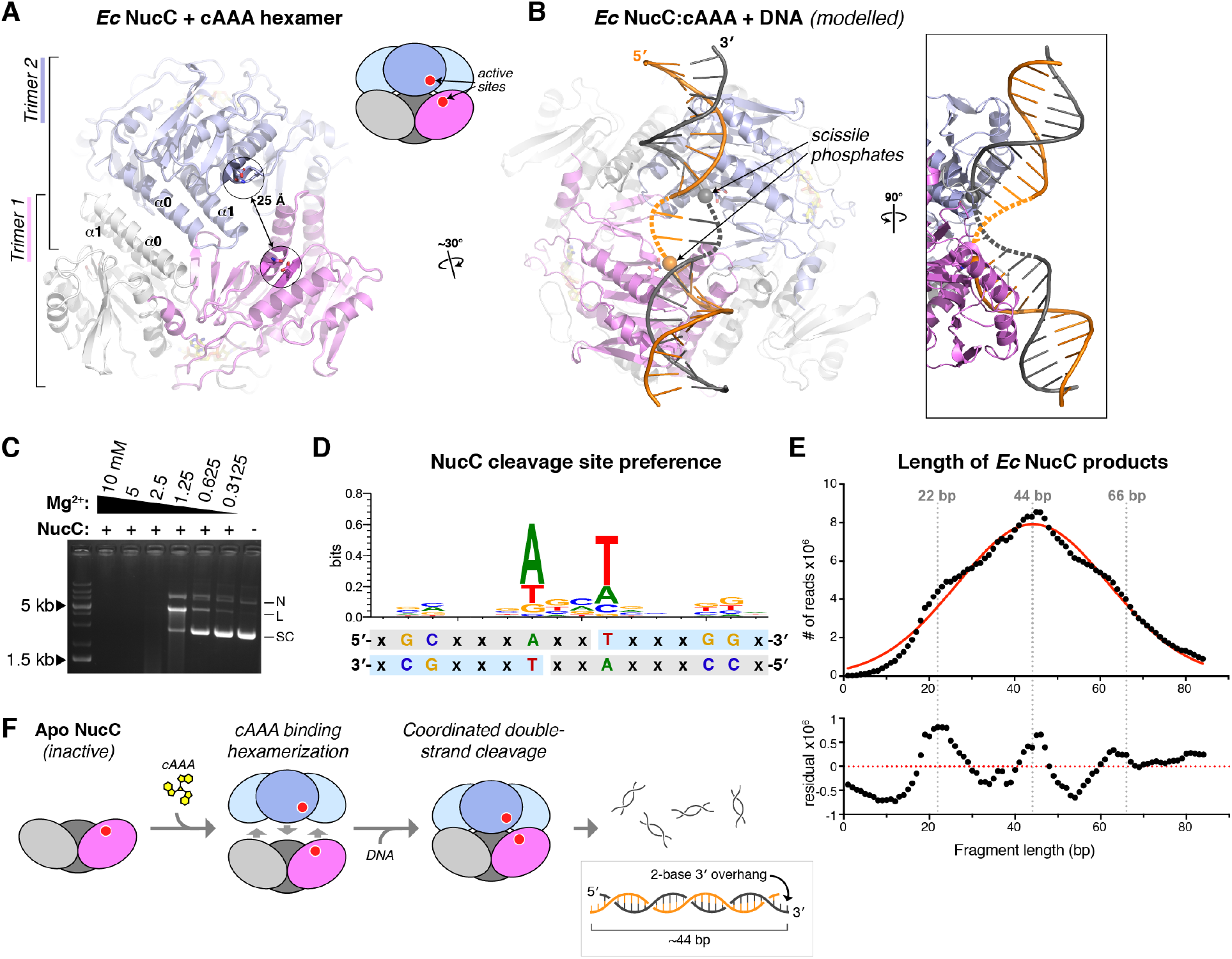
NucC generates double-strand cuts with two-base 3’ overhangs. **(A)** View of the cAAA-bound *Ec* NucC hexamer, showing the juxtaposition of two active sites ~25 Å apart. **(B)** Model of a NucC hexamer bound to DNA, assembled from separately overlaying the NucC monomers colored violet and blue with a structure of SgrAI bound to DNA (PDB ID 3DW9; (Dunten et al., 2008)). The two modelled DNAs can be joined by a bent-two base-pair linker (dotted lines) to generate a contiguous dsDNA (two strands colored gray and orange). Cleavage of the scissile phosphates (spheres) at each active site would generate a double-strand break with a two-base 3’ overhang. **(C)** Plasmid cleavage by *Ec* NucC in limiting Mg^2+^. Supercoiled DNA (SC) is converted to both nicked (N) and linear (L) forms, demonstrating significant double-strand cleavage. **(D)** Analysis of NucC DNA cleavage site preference, from mapping ~600,000 cleavage sites from fully-digested plasmid DNA. Light-blue indicates the sequenced strand; the two-fold symmetry of the cleavage site supports a model of double-strand cleavage resulting in two-base 3’ overhangs. **(E)** Length distribution of *Ec* NucC products after complete digestion, using high-throughput sequencing read lengths from a 343 million-read dataset. Red line shows a Gaussian fit, and bottom panel shows residual values from the Gaussian fit. **(F)** Model for hexamerization and coordinated double-strand cleavage by NucC. Inactive NucC trimers associate upon binding cAAA to form active hexamers. Coordinated cleavage at pairs of NucC active sites juxtaposed by hexamerization results in double-strand breaks with two-base 3’ overhangs.

To determine whether NucC generates single-or double-strand breaks, we used limiting Mg^2+^ conditions to enrich for singly-cut plasmid in a plasmid digestion assay (Fig. 3C). While NucC did generate a nicked species in this assay, we also observed a significant fraction of linear DNA that could only arise through cleavage of both DNA strands at a single site. To further support the idea that NucC can generate double-strand breaks, we next completely digested a plasmid with NucC and deep-sequenced the resulting short fragments. To avoid losing information about DNA ends by “blunting” during sequencing library preparation, we denatured the fragments and sequenced individual DNA strands (see **Materials and Methods**). We thereby mapped ~1 million cleavage events, enabling us to assemble a picture of any preferred sequence motifs at NucC cleavage sites. We observed a striking two-fold symmetry in the preferred cleavage sites, with a strong preference for “T” at the 5’ end (+1 base) of the cleaved fragment, and an equally strong preference for “A” at the −3 position (Fig. 3D). Similarly, we observed a weak preference for “G” at the +5 and +6 bases, and for “C” at the −7 and −8 bases. The two-fold symmetry of preferred NucC cleavage sites strongly suggests that two NucC active sites cooperate to cleave both strands of DNA. Moreover, the location of the breaks on each strand (as judged by the first sequenced base of the fragments) suggests that double-stranded cleavage by NucC results in two-base 3’ overhangs, consistent with our structural modeling (Fig. 3D and Fig. 3F).

We used the same sequencing dataset to precisely measure the length distribution of NucC products. NucC product sizes followed a roughly Gaussian distribution with a mean of ~44 bases, in agreement with our electrophoretic results (Fig. 3E). Curiously, the size distribution showed a systematic variation from a Gaussian distribution, with slight enrichment of product lengths around 22, 44, and 66 base pairs in length. These data suggest that in addition to coordinated DNA cleavage by pairs of NucC active sites, cleavage by the three pairs of active sites in a NucC hexamer may be weakly coordinated, perhaps through wrapping of a single DNA around the complex to contact multiple sets of active sites.

### Type III CRISPR systems encode NucC homologs

While exploring the distribution of NucC homologs in bacteria, we identified a group of NucC homologs with ~50% sequence identity to *Ec* NucC and *Pa* NucC, that are encoded not within CD-NTase operons but rather within type III CRISPR systems (Fig. 4A-C). Type III CRISPR systems encode a multi-subunit effector complex, that when activated by target binding synthesizes a range of cyclic oligoadenylate second messengers with 2-6 bases (cA_2_-cA_6_) (Kazlauskiene et al., 2017; Niewoehner et al., 2017). These second messengers have been shown to activate accessory RNases in the CRISPR operons, with Csm6 activated by cA_6_ and Csx1 activated by cA_4_ (Han et al., 2018; Kazlauskiene et al., 2017; Niewoehner et al., 2017). The presence of NucC homologs in type III CRISPR systems suggests that these enzymes may function similarly to their relatives in CD-NTase operons, with activation dependent on cAAA/cA_3_ synthesized by the CRISPR effector complexes.

**Figure 4.**
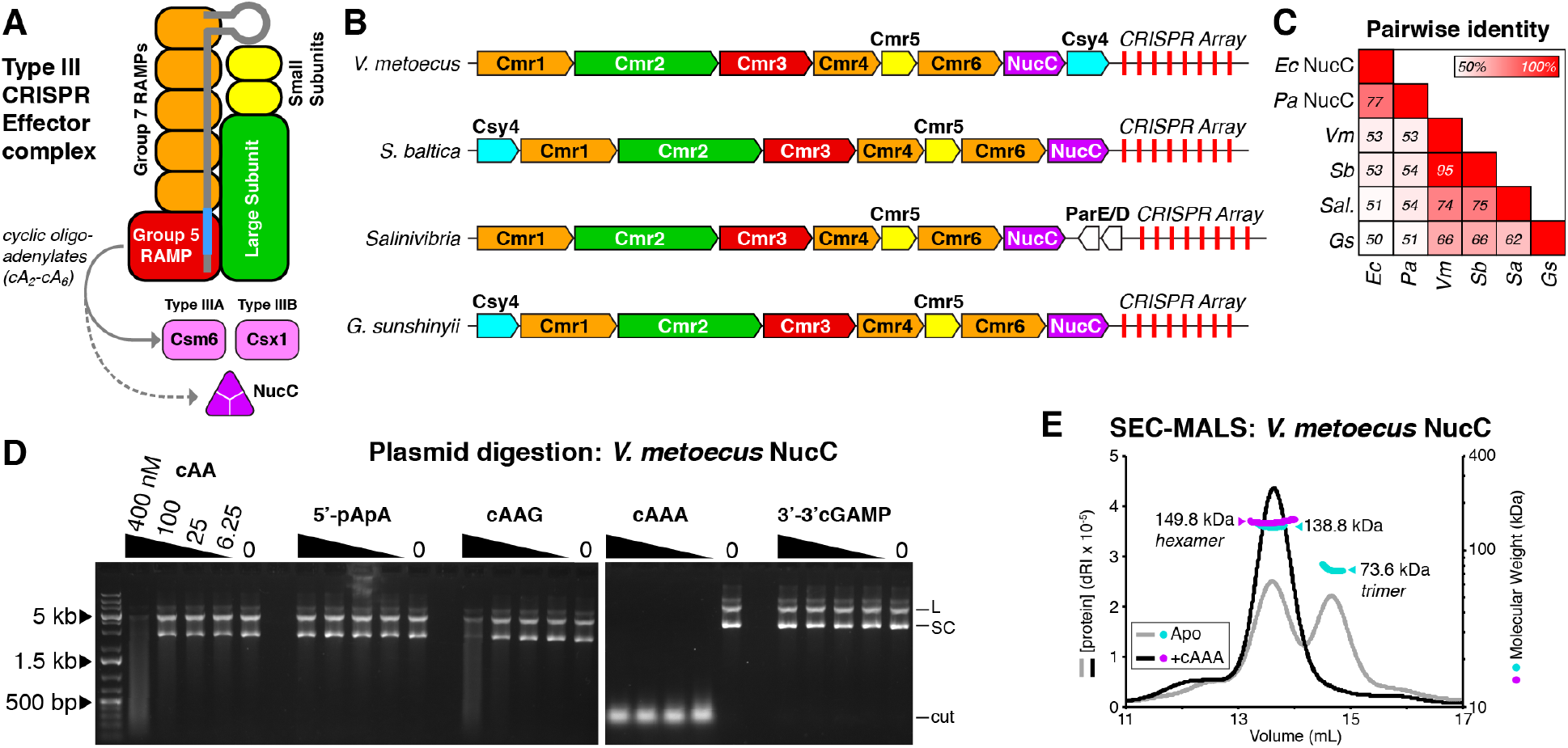
NucC is an accessory nuclease in Type III CRISPR systems. **(A)** Schematic of Type III CRISPR effector complexes (adapted from (Makarova et al., 2017)). The Group 5 RAMP subunit (red) produces cyclic oligoadenylates that in some type III systems activate an accessory RNase, Csm6 or Csx1. **(B)** Schematic of four bacterial operons encoding type III CRISPR systems with NucC homologs (*V. metoecus* = *Vibrio metoecus* RC341; *S. baltica* = *Shewanella baltica* OS625; *Salinivibria* = *Salinivibria* sp. MA351; *G. sunshinyii* = *Gynuella sunshinyii* YC6258). **(C)** Pairwise sequence identity of *Ec* NucC, *Pa* NucC, and four type III CRISPR-associated NucC homologs (*Vm* = *Vibrio metoecus* RC341; *S. baltica* = *Sb*OS625; *Sal.* = *Salinivibria* sp. MA351; *Gs* = *Gynuella sunshinyii* YC6258). **(D)** Plasmid digestion assay with *V. metoecus* NucC in the presence of nucleotide-based second messengers. **(E)** SEC-MALS of *V. metoecus* NucC in either the Apo state (gray/cyan) or after pre-incubation with cAAA (black/magenta). Calculated MW of the NucC homotrimer = 84.1 kDa; homohexamer = 168.1 kDa.

We cloned several CRISPR-associated NucC proteins, and successfully purified one such enzyme from *Vibrio metoecus*, a close relative of *V. cholerae* (Kirchberger et al., 2014). We found that *V. metoecus* NucC is a DNA endonuclease that is strongly activated by cAAA, and weakly activated by both cAA and cAAG, mirroring our findings with *Ec* NucC (Fig. 4D) (Ye et al., 2019). By SEC-MALS, we found that *V. metoecus* NucC forms a mix of trimers and hexamers on its own, and forms solely hexamers after pre-incubation with cAAA (Fig. 4E). These data strongly support the idea that CRISPR-associated NucC homologs share a common activation and DNA cleavage mechanism with their relatives in CD-NTase operons. Thus, NucC homologs have been incorporated into type III CRISPR systems, where they are likely activated upon target recognition and cAAA synthesis by these systems’ effector complexes.

## Discussion

Bacteria possess an extraordinary variety of pathways to defend themselves against foreign threats including bacteriophages, invasive DNA elements like plasmids, and other bacteria. A large class of defense pathways, including classical restriction-modification systems and CRISPR-Cas systems, specifically defend the bacterial genome through the targeted action of DNA and RNA nucleases. Here, we describe the structure and mechanism of a restriction endonuclease-like protein, NucC, that is activated by a cyclic trinucleotide second messenger, cAAA. We have recently shown that the intact NucC-containing operon from a patient-derived *E. coli* strain confers immunity to bacteriophage λ infection, and that both cAAA synthesis by this operon’s CD-NTase and the DNA cleavage activity of NucC are required for immunity (Ye et al., 2019). Thus, NucC is part of a widespread bacterial defense pathway that senses and responds to foreign threats including bacteriophages.

While evolutionarily related to restriction endonucleases, NucC adopts a unique homotrimeric structure with an allosteric binding pocket for its cyclic trinucleotide activator. This pocket is assembled from embellishments to the core nuclease fold, which include a “gate loop” that closes over the bound second messenger. cAAA binding stimulates assembly of two NucC trimers into a hexamer, juxtaposing pairs of active sites and activating double-strand DNA cleavage. While the nuclease does possess weak sequence preference, it is largely nonspecific and cleaves DNA to an average length of ~50 base pairs. NucC is apparently insensitive to DNA methylation (Ye et al., 2019), meaning the bacterial genome may not be protected from NucC cleavage. Thus, we propose that CD-NTase operons encoding NucC may function as “abortive infection” pathways, in which an infected cell commits suicide to avoid producing more bacteriophage. Alternatively, localized activation of NucC within an infected cell may result in specific phage genome destruction.

Bacterial operons encoding CD-NTases are widespread among environmental and pathogenic bacteria, but are sparsely distributed and often associated with transposase or integrase genes and/or conjugation systems, suggesting that they are mobile DNA elements that likely confer some selective advantage on their bacterial host (Burroughs et al., 2015; Whiteley et al., 2019). These operons encode diverse putative effector proteins including proteases, exonucleases, and two families of putative DNA endonucleases (Burroughs et al., 2015). Given the structure of these operons, it is likely that these effectors are specifically activated by the second messenger synthesized by their associated CD-NTase (Whiteley et al., 2019). Bacterial CD-NTases have been classified into eight major clades, and while members of several clades have been shown to synthesize specific cyclic di- and trinucleotides (Whiteley et al., 2019), the full spectrum of products synthesized by these enzymes remains an important question. Given our data on NucC structure and activation, we propose that all bacterial CD-NTases associated with NucC – which includes members of clades C01 (including *E. coli* MS115-1 CdnC), C03, and D05 (including *P. aeruginosa* ATCC27853 CdnD) (Whiteley et al., 2019) – likely synthesize cAAA. While CD-NTases in clades C01 and C03 are primarily associated with NucC effectors, clade D05 CD-NTases are associated with a range of effectors (Whiteley et al., 2019). One key question is whether all of these effectors are activated by cAAA, or whether their associated CD-NTases synthesize distinct second messengers. More broadly, a full accounting of bacterial CD-NTase products, and of the effector proteins they activate, remains an important area for future study.

While NucC is primarily found within CD-NTase operons, we also identify a group of type III CRISPR-Cas systems that encode homologs of NucC. Type III CRISPR systems uniquely encode nonspecific accessory RNases, Csm6 and Csx1, that are activated by cyclic oligoadenylates synthesized by their effector complexes (Han et al., 2018; Kazlauskiene et al., 2017; Niewoehner et al., 2017). We show that at least one type III CRISPR-associated NucC homolog from *Vibrio metoecus* is a cAAA-stimulated DNA endonuclease that forms hexamers in the presence of cAAA. Thus, the production of a range of cyclic oligoadenylates including cAAA by type III CRISPR effector complexes has allowed NucC to be integrated into these systems as a tightly-controlled accessory nuclease. Given the high probability that other CD-NTase associated effectors are activated by cyclic oligoadenylates, it is possible that additional cases of such integration remain to be identified.

## Materials and Methods

### Protein Expression, Purification, and Characterization

Full length NucC genes from *E. coli* MS115-1 (synthesized by Invitrogen/Geneart), *P. aeruginosa* ATCC 27853 (amplified from genomic DNA), *Vibrio metoecus* sp. RC341 (synthesized by Invitrogen/Geneart), and *Gynuella sunshinyii* YC6258 (synthesized by Invitrogen/Geneart) were cloned into UC Berkeley Macrolab vector 1B (Addgene #29653) to generate N-terminal fusions to a TEV protease-cleavable His_6_-tag. Mutants of *Ec* NucC were generated by PCR mutagenesis.

Proteins were expressed in *E. coli* strain Rosetta2 pLysS (EMD Millipore) by induction with 0.25 mM IPTG at 20°C for 16 hours. For selenomethionine derivatization of *Ec* NucC, cells were grown in M9 minimal medium, then supplemented with amino acids (Leu, Ile and Val (50 mg/L), Phe, Lys, Thr (100 mg/L) and Selenomethionine (60 mg/L)) upon induction with IPTG.

For protein purification, cells were harvested by centrifugation, suspended in resuspension buffer (20 mM Tris-HCl pH 8.0, 300 mM NaCl, 10 mM imidazole, 1 mM dithiothreitol (DTT) and 10% glycerol) and lysed by sonication. Lysates were clarified by centrifugation (16,000 rpm 30 min), then supernatant was loaded onto a Ni^2+^ affinity column (HisTrap HP, GE Life Sciences) pre-equilibrated with resuspension buffer. The column was washed with buffer containing 20 mM imidazole and 100 mM NaCl, and eluted with a buffer containing 250 mM imidazole and 100 mM NaCl. The elution was loaded onto an anion-exchange column (Hitrap Q HP, GE Life Sciences) and eluted using a 100-600 mM NaCl gradient. Fractions containing the protein were pooled and mixed with TEV protease (1:20 protease:NucC by weight) plus an additional 100 mM NaCl, then incubated 16 hours at 4°C for tag cleavage. Cleavage reactions were passed over a Ni^2+^ affinity column, and the flow-through containing cleaved protein was collected and concentrated to 2 mL by ultrafiltration (Amicon Ultra-15, EMD Millipore), then passed over a size exclusion column (HiLoad Superdex 200 PG, GE Life Sciences) in a buffer containing 20 mM Tris-HCl pH 8.0, 300 mM NaCl, and 1 mM DTT. Purified proteins were concentrated by ultrafiltration and stored at 4°C for crystallization, or aliquoted and frozen at −80°C for biochemical assays. All mutant proteins were purified as wild-type.

For characterization of oligomeric state by size exclusion chromatography coupled to multi-angle light scattering (SEC-MALS), 100 μL of purified protein/complex at 2-5 mg/mL was injected onto a Superdex 200 Increase 10/300 GL column (GE Life Sciences) in a buffer containing 20 mM HEPES-NaOH pH 7.5, 300 mM NaCl, 5% glycerol, and 1 mM DTT. Light scattering and refractive index profiles were collected by miniDAWN TREOS and Optilab T-rEX detectors (Wyatt Technology), respectively, and molecular weight was calculated using ASTRA v. 6 software (Wyatt Technology).

### Second messenger molecules

Cyclic and linear dinucleotides, including 5’-pApA, 3’,3’-cGAMP, and 3’,3’-cyclic di-AMP, were purchased from Invivogen. Cyclic trinucleotides were generated enzymatically by *Ec*-CdnD02 from *Enterobacter cloacae*, which was cloned and purified as described (Whiteley et al., 2019). Briefly, a 40 mL synthesis reaction contained 500 nM *Ec*-CdnD02 and 0.25 mM each ATP and GTP (ATP alone for large-scale synthesis of cAAA) in reaction buffer with 12.5 mM NaCl, 20 mM MgCl_2_, 1 mM DTT, and 10 mM Tris-HCl pH 9.0. The reaction was incubated at 37°C for 16 hours, then 2.5 units/mL reaction volume (100 units for 40 mL reaction) calf intestinal phosphatase was added and the reaction incubated a further 4 hours at 37°C. The reaction was heated to 65°C for 30 minutes, and centrifuged 20 minutes at 4,000 RPM to remove precipitated protein. Reaction products were separated by ion-exchange chromatography (Mono-Q, GE Life Sciences) using a gradient from 0 to 2 M ammonium acetate. Three major product peaks (Fig. S2A) were pooled, evaporated using a speedvac, then resuspended in water and analyzed by LC MS/MS. cAAA synthesized by the *E. coli* MS115-1 CdnC:HORMA complex was similarly purified and verified to activate NucC equivalently to that synthesized by *Ec*-CdnD02 (Ye et al., 2019).

### Mass spectrometry

Whiteley et al. previously identified the major product of *Ec*-CdnD02 as cAAG by NMR (Whiteley et al., 2019). To further characterize the products of *Ec*-CdnD02, we performed liquid chromatography-tandem mass spectrometry (LC-MS/MS) (Fig. S2B-D). LC-MS/MS analysis was performed using a Thermo Scientific Vanquish UHPLC coupled to a Thermo Scientific Q Exactive™ HF Hybrid Quadrupole-Orbitrap™ Mass Spectrometer, utilizing a ZIC-pHILIC polymeric column (100 mm × 2.1 mm, 5 μm) (EMD Millipore) maintained at 45 °C and flowing at 0.4 mL/min. Separation of cyclic trinucleotide isolates was achieved by injecting 2 μL of prepared sample onto the column and eluting using the following linear gradient: (A) 20 mM ammonium bicarbonate in water, pH 9.6, and (B) acetonitrile; 90% B for 0.25 minutes, followed by a linear gradient to 55% B at 4 minutes, sustained until 6 minutes. The column was re-equilibrated for 2.50 minutes at 90% B.

Detection was performed in positive ionization mode using a heated electrospray ionization source (HESI) under the following parameters: spray voltage of 3.5 kV; sheath and auxiliary gas flow rate of 40 and 20 arbitrary units, respectively; sweep gas flow rate of 2 arbitrary units; capillary temperature of 275 °C; auxiliary gas heater temperature of 350°C. Profile MS1 spectra were acquired with the following settings; mass resolution of 35,000, AGC volume of 1×106, maximum IT of 75 ms, with a scan range of 475 to 1050 m/z to include z=1 and z=2 ions of cyclic trinucleotides. Data dependent MS/MS spectra acquisition was performed using collision-induced dissociation (CID) with the following settings: mass resolution of 17,500; AGC volume of 1×105; maximum IT of 50 ms; a loop count of 5; isolation window of 1.0 m/z; normalized collision energy of 25 eV; dynamic exclusion was not used. Data reported is for the z=1 acquisition for each indicated cyclic trinucleotide.

### Isothermal Titration Calorimetry

Isothermal titration calorimetry was performed at 20°C on a Microcal ITC 200 (Malvern Panalytical) in a buffer containing 20 mM Tris-HCl (pH 8.5), 200 mM NaCl, 5 mM MgCl_2_, 1 mM EDTA, and 1 mM TCEP. Nucleotides at 1 mM were injected into an analysis cell containing 100 μM *Ec* NucC (Fig. S3A-G, Table S2). A second round of ITC was performed with 5’pApA and 3’,3’c-di-AMP in a buffer containing 10 mM Tris-HCl (pH 7.5), 25 mM NaCl, 10 mM MgCl_2_, and 1 mM TCEP (Fig. S3H-K, Table S3).

### Crystallization and structure determination

#### E. coli NucC

We obtained crystals of *E. coli* NucC in the Apo state in hanging drop format by mixing protein (8-10 mg/mL) in crystallization buffer 1:1 with well solution containing 27% PEG 400, 400 mM MgCl_2_, and 100 mM HEPES pH 7.5 (plus 1 mM TCEP for selenomethionine-derivatized protein). For cryoprotection, we supplemented PEG 400 to 30%, then flash-froze crystals in liquid nitrogen and collected diffraction data on beamline 14-1 at the Stanford Synchrotron Radiation Lightsource (support statement below). We processed all datasets with the SSRL autoxds script, which uses XDS (Kabsch, 2010) for data indexing and reduction, AIMLESS (Evans and Murshudov, 2013) for scaling, and TRUNCATE (Evans, 2006) for conversion to structure factors. We determined the structure by single-wavelength anomalous diffraction methods with a 1.82 Å resolution dataset from selenomethione-derivatized protein. We identified heavy-atom sites using hkl2map (Pape et al., 2004) (implementing SHELXC and SHELXD (Sheldrick, 2010)), then provided those sites to the PHENIX Autosol wizard (Terwilliger et al., 2009), which uses PHASER (McCoy et al., 2007) for phase calculation and RESOLVE for density modification (Terwilliger, 2003a) and automated model building (Terwilliger, 2003b). We manually rebuilt the model in COOT (Emsley et al., 2010), followed by refinement in phenix.refine (Afonine et al., 2012) using positional, individual B-factor, and TLS refinement (statistics in Table S1).

We obtained crystals of *E. coli* NucC bound to 5’-pApA or cAAA in hanging drop format by mixing protein (8-10 mg/mL) in crystallization buffer plus 0.1 mM 5’-pApA (Invivogen) or cAAA 1:1 with well solution containing 17-24% PEG 3350, 0.1 M Na/K tartrate, and 25 mM Tris-HCl pH 8.0. For cryoprotection, we added 10% glycerol, then flash-froze crystals in liquid nitrogen and collected diffraction data on beamline 24ID-E at the Advanced Photon Source at Argonne National Lab (support statement below). We processed datasets with the RAPD data-processing pipeline, which uses XDS, AIMLESS, and TRUNCATE as above. We determined the structures by molecular replacement using the structure of Apo NucC, manually rebuilt the model in COOT, and refined in phenix.refine using positional, individual B-factor, and TLS refinement (statistics in Table S1). For both 5’-pApA and cAAA, ligands were fully built and assigned an occupancy of 0.33 to account for their location on a three-fold crystallographic rotation axis. The ligands were assigned a “custom-nonbonded-symmetry-exclusion” in phenix.refine to allow symmetry overlaps during refinement.

#### P. aeruginosa NucC

We obtained crystals of *P. aeruginosa* ATCC27853 NucC bound to cAAA by mixing protein (12 mg/mL) in cerystallization buffer plus 0.1 mM cAAA 1:1 with well solution containing 0.1 M CHES pH 9.9, 0.2 M ammonium acetate, and 20% PEG 3350. Crystals were cryoprotected with an additional 10% PEG 400, flash-frozen in liquid nitrogen, and a 2.6 Å diffraction dataset was collected at beamline 24ID-E at the Advanced Photon Source. We manually processed the dataset with XDS and TRUNCATE, and determined the structure by molecular replacement in PHASER using the structure of cAAA-bound *Ec* NucC as a search model. We manually rebuilt the model in COOT, and refined in phenix.refine using positional, individual B-factor, and TLS refinement (statistics in Table S1).

### SSRL Support Statement

Use of the Stanford Synchrotron Radiation Lightsource, SLAC National Accelerator Laboratory, is supported by the U.S. Department of Energy, Office of Science, Office of Basic Energy Sciences under Contract No. DE-AC02-76SF00515. The SSRL Structural Molecular Biology Program is supported by the DOE Office of Biological and Environmental Research, and by the National Institutes of Health, National Institute of General Medical Sciences (including P41GM103393). The contents of this publication are solely the responsibility of the authors and do not necessarily represent the official views of NIGMS or NIH.

### APS NE-CAT Support Statement

This work is based upon research conducted at the Northeastern Collaborative Access Team beamlines, which are funded by the National Institute of General Medical Sciences from the National Institutes of Health (P41 GM103403). The Eiger 16M detector on 24-ID-E beam line is funded by a NIH-ORIP HEI grant (S10OD021527). This research used resources of the Advanced Photon Source, a U.S. Department of Energy (DOE) Office of Science User Facility operated for the DOE Office of Science by Argonne National Laboratory under Contract No. DE-AC02-06CH11357.

### Nuclease assays

For all nuclease assays, UC Berkeley Macrolab plasmid 2AT (Addgene #29665; 4,731 bp; sequence in Supplemental Information) was used. *Ec* NucC (10 nM unless otherwise indicated) and second messenger molecules (10 nM unless otherwise indicated) were mixed with 1 μg plasmid DNA in a buffer containing 10 mM Tris-HCl (pH 7.5), 25 mM NaCl, 10 mM MgCl_2_, and 1 mM DTT (50 μL reaction volume), incubated 10 min at 37°C, then separated on a 1.2% agarose gel. Gels were stained with ethidium bromide and imaged by UV illumination.

### High-throughput sequencing and data analysis

For high-throughput analysis of NucC products, 1 μg of vector 2AT was digested to completion by *Ec* NucC, separated by agarose gel, then the prominent ~50 bp band was removed from the gel and gel-purified (Qiagen Minelute). 80 ng of purified DNA was prepared for sequencing using the Swift Biosciences Accel-NGS 1S Plus DNA Library kit, then sequenced on an Illumina HiSeq 4000 (paired-end, 100 base reads), yielding ~402 million read pairs. The Accel-NGS 1S kit: (1) separates DNA fragments into single strands; (2) ligates the Adapter 1 oligo to the 3’ end of the single-stranded fragment using an “adaptase” step that adds ~8 bp of low-complexity sequence to the fragment before adapter ligation; (3) ligates the Adapter 2 oligo to the 5’ end of the original fragment. After sequencing, read 1 (R1) represents the 5’ end of the original fragment, and read 2 (r2) represents the 3’ end (preceded by ~ 8 bp of low-complexity sequence resulting from the adaptase step).

For NucC fragment length analysis, we took advantage of the fact that most NucC products are less than 100 base pairs in length. We identified the (reverse complement of the) 3’ end of Adapter 1 (full sequence: 5’AATGATACGGCGACCACCGAGATCTACACTCTTTCCCTACAC GACGCTCT<u>TCCGATCT</u> 3’) in R1, searching for the sequence “AGATCGGA”. We calculated the length of each NucC fragment as: (length of R1 prior to AGATCGGA) minus 8 (average length of adaptase-added low-complexity sequence). The graph in Fig. 4C represents the lengths of 343 million NucC fragments.

For NucC cleavage site identification (Fig. 4B), we truncated a 1-million read subset of R1 to 15 bases, then mapped to the 2AT plasmid sequence (with ends lengthened by 100 bases to account for circularity of the plasmid) using bowtie (Langmead et al., 2009) with the command: “bowtie -a -n 0 -m 3 -best -t”. Of the successfully-mapped reads (594,183 of 1 million, 59.4%), 288,609 reads mapped to the forward strand and 305,574 mapped to the reverse strand. We extracted 20 bp of sequence surrounding each mapped 5’ end (first base of R1), and input the resulting alignment of 594,183 sequences into WebLogo 3 (http://weblogo.threeplusone.com) (Crooks et al., 2004).

## End Matter

### Author Contributions and Notes

Conceptualization, RKL, QY, KDC; Methodology, RKL, QY, LP, ITM, JDW, ATW, BL, MJ, PJK, KDC; Investigation, RKL, QY, LP, KRB, ITM, JDW, KDC; Writing – Original Draft, RKL, QY, KDC; Writing – Review & Editing, RKL, QY, MJ, PJK, KDC; Visualization, RKL, QY, ITM, JDW, KDC; Supervision, MJ, JJM, PJK, KDC; Project Administration, KDC; Funding Acquisition, MJ, JJM, PJK, KDC.

The authors declare no conflict of interest.

## Acknowledgments

The authors thank the staffs of the Stanford Synchrotron Radiation Lightsource and the Advanced Photon Source NE-CAT beamlines for assistance with crystallographic data collection; A. Bobkov (Sanford Burnham Prebys Medical Discovery Institute, Protein Analysis Core) for assistance with isothermal titration calorimetry; and A. Desai, M. Daugherty, and J. Pogliano for critical reading and helpful suggestions. RKL was supported by the UC San Diego Quantitative and Integrative Biology training grant (NIH T32 GM127235). ITM was supported by NIH T32 HL007444, T32 GM007752 and F31 CA236405. JDW was supported by NIH K01 DK116917 and P30 DK063491. MJ was supported by NIH S10 OD020025 and R01 ES027595. ATW was supported as a fellow of The Jane Coffin Childs Memorial Fund for Medical Research. BL was supported as a Herchel Smith Graduate Research Fellow. JJM was supported by NIH/NIAID R01AI018045 and R01AI026289. PJK was supported from the Richard and Susan Smith Foundation and the Parker Institute for Cancer Immunotherapy. KDC was supported from the Ludwig Institute for Cancer Research and the University of California, San Diego.

---

**Figure S1.**
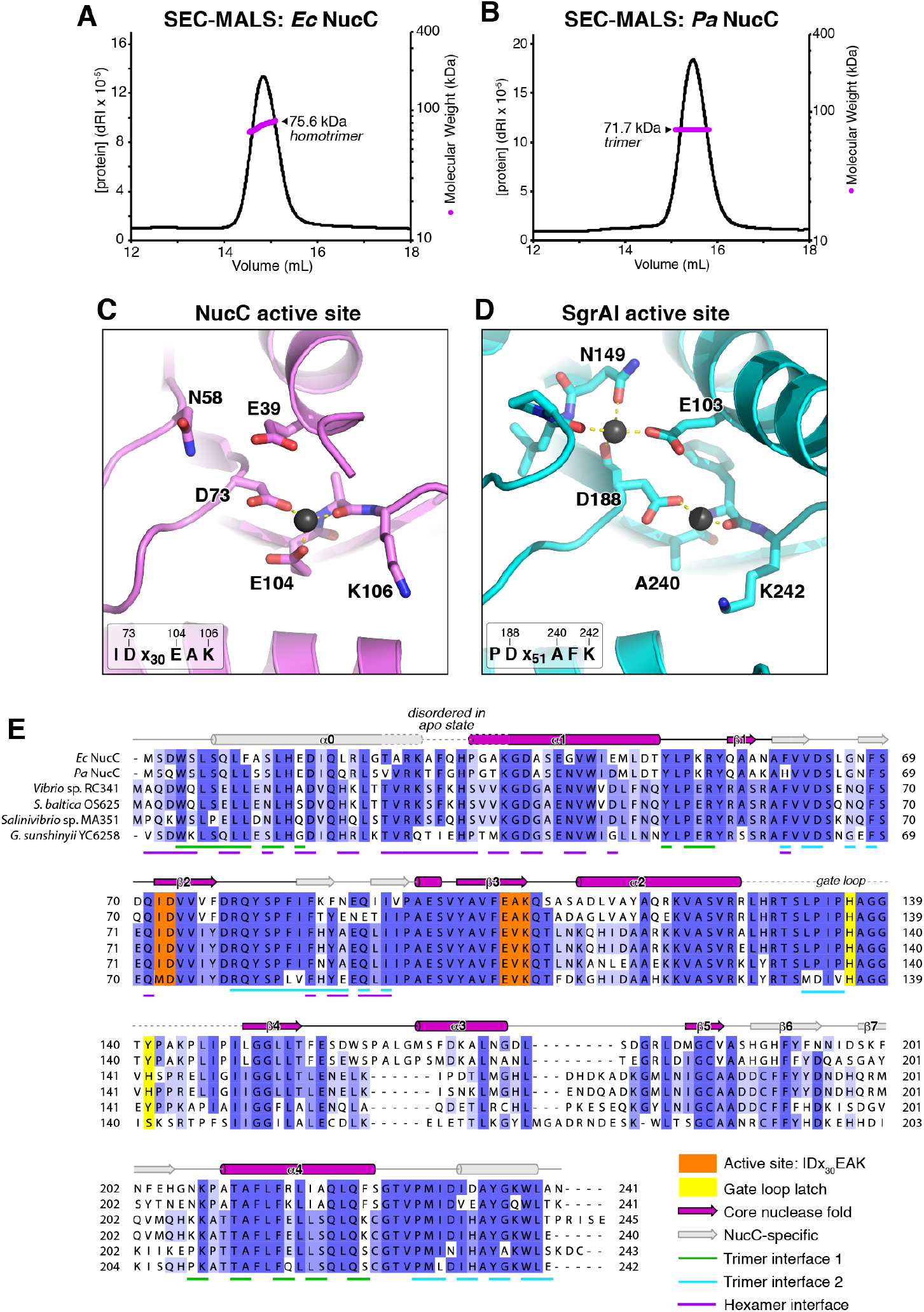
NucC sequence and structure. **(A)** Size exclusion chromatography coupled to multi-angle light scattering (SEC-MALS) of purified *Ec* NucC (theoretical trimer = 80.1 kDa). **(B)** SEC-MALS of purified *Pa* NucC (theoretical trimer = 80.0 kDa). **(C)** Closeup view of the NucC active site, with conserved active-site residues shown as sticks and Mg^2+^ shown as black sphere. *Inset* shows active-site consensus motif. **(D)** Closeup view of the active site of restriction endonuclease SgrAI, from a DNA-bound structure (PDB ID 3DW9) (Dunten et al., 2008). Bound DNA, which coordinates one Mn^2+^ ion with D188 and the main-chain carbonyl group of F241, is not shown. *Inset* shows active-site consensus motif. **(E)** Sequence alignment of *E. coli* MS115-1 (*Ec*) and *P. aeruginosa* ATCC27853 (*Pa*) NucC with four NucC homologs associated with Type IIIB CRISPR systems (Fig. 4; all sequences in Table S4). Secondary structure shown is that of *Ec* NucC, with elements of the core nuclease fold in violet and embellishments in gray (Fig. 1B). Orange highlights indicate the active-site consensus motif (IDx_30_EAK) and yellow highlights indicate the gate loop latch residues H136 and Y141. Green and cyan underlines indicate residues involved in the two interfaces that mediate NucC trimer assembly, calculated by PISA (Krissinel and Henrick, 2007).

**Figure S2.**
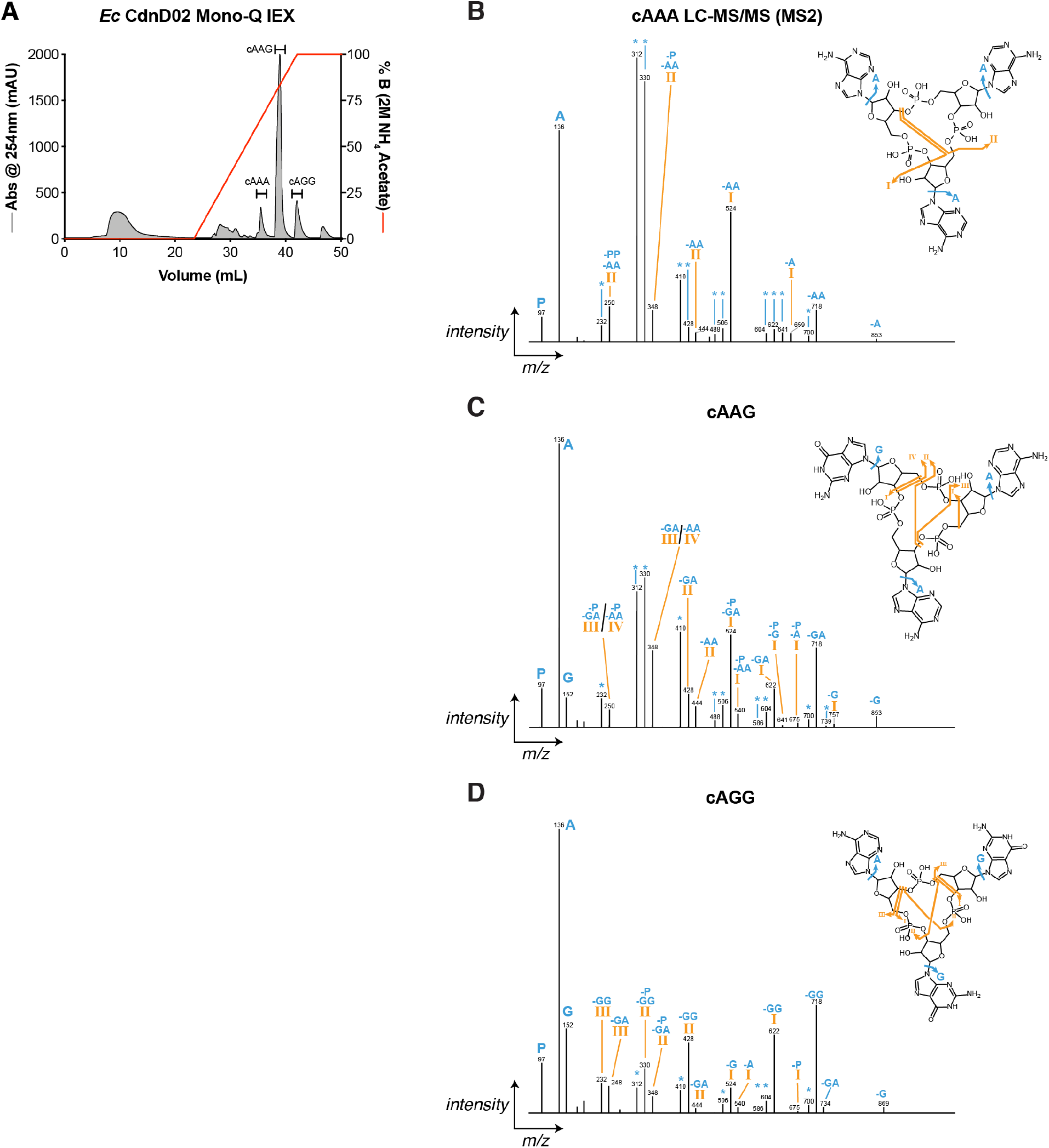
Cyclic trinucleotide synthesis by *Enterobacter cloacae* CdnD02. **(A)** Mono-Q ion-exchange chromatography elution profile of reaction products generated by *Enterobacter cloacae* CdnD02 when supplied with GTP and ATP. The most prominent peak (cAAG) was previously characterized by NMR (Whiteley et al., 2019). The cAAG peak was verified, and two minor products identified as cAAA and cAGG, by LC-MS/MS mass spectrometry (panels B-D). **(B-D)** Mass fragmentation pattern of Mono-Q peak 1 (cAAA, panel B), peak 2 (cAAG, panel C), and peak 3 (cAGG, panel D). Roman numerals indicate associated bond cleavage, with asterisk (*) indicating additional loss of water. “-A”, “-G”, and “-P” indicate loss of adenine, guanine, and phosphate, respectively.

**Figure S3.**
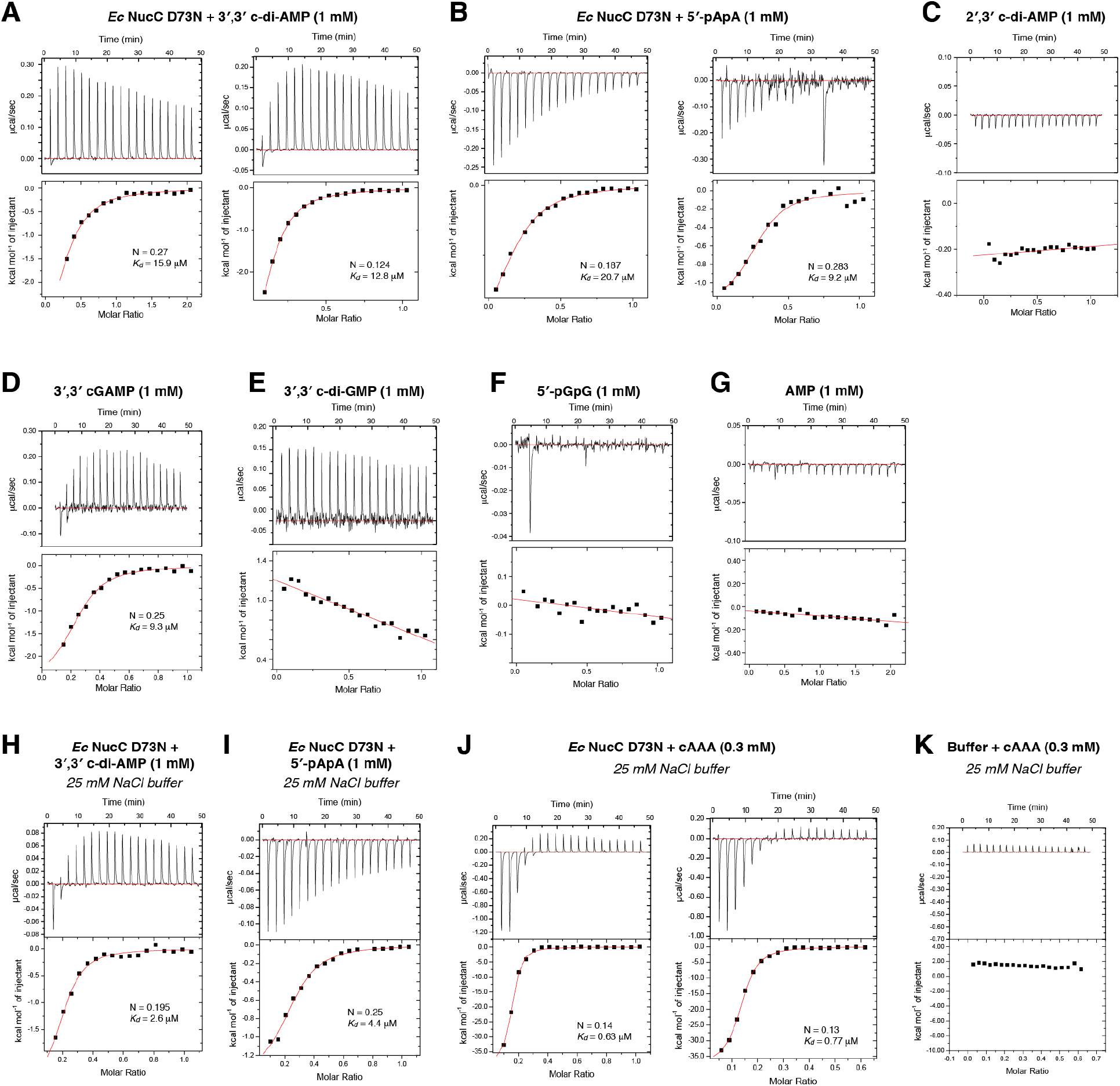
Binding of NucC to second messengers. **(A)** Binding of *Ec* NucC D73N to 3’,3’ cyclic di-AMP in high-salt buffer by isothermal titration calorimetry. See Table S2 for affinity and stoichiometry of binding for panels A-G. **(B)** Binding of *Ec* NucC D73N to 5’-pApA (linear di-AMP). **(C)** Binding of *Ec* NucC D73N to 2’,3’ cyclic di-AMP. **(D)** Binding of *Ec* NucC D73N to 3’,3’ cyclic GMP-AMP (cGAMP). **(E)** Binding of *Ec* NucC D73N to 3’,3’ cyclic di-GMP. **(F)** Binding of *Ec* NucC D73N to 5’-pGpG (linear di-GMP). **(G)** Binding of *Ec* NucC D73N to AMP. **(H)** Binding of *Ec* NucC D73N to 3’,3’ cyclic di-AMP in low-salt buffer. See Table S3 for affinity and stoichiometry of binding for panels H-J. **(I)** Binding of *Ec* NucC D73N to 5’-pApA (linear di-AMP) in low-salt buffer. **(J)** Binding of *Ec* NucC D73N to cAAA in low-salt buffer. **(K)** Negative control titration of low-salt buffer with cAAA.

**Figure S4.**
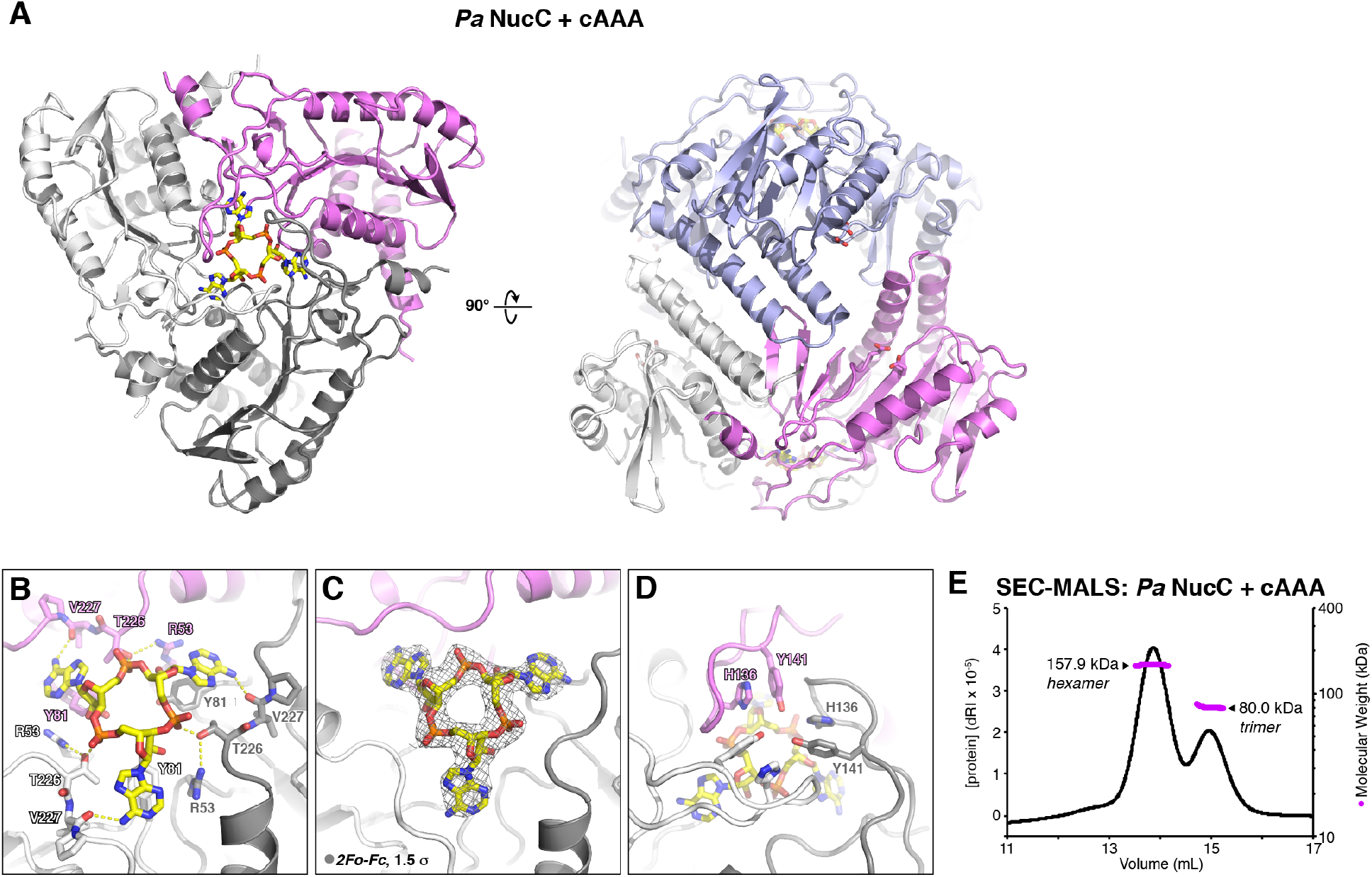
Structure of *Pa* NucC. **(A)** *Left:* Bottom view of a *Pa* NucC trimer (chains colored violet, gray, and white) with bound cAAA shown as sticks. The *Pa* NucC trimer overlays with the cAAA-bound *Ec* NucC trimer with an overall Cα r.m.s.d. of 0.51 Å (649 Cα pairs). *Right:* Side view of a *Pa* NucC hexamer in the cAAA-bound state (one trimer colored as in panel A, and the second trimer colored light blue). Active site residues Asp73 and Glu104 are shown as sticks. **(B)** Closeup view of cAAA binding to *Pa* NucC. **(C)** *2Fo-Fc* electron density for cAAA at 2.6 Å resolution, contoured at 1.5 σ. **(D)** View of the *Pa* NucC gate loops enclosing the bound cAAA. **(E)** Size exclusion chromatography coupled to multi-angle light scattering (SEC-MALS) of purified *Pa* NucC after pre-incubation with cAAA. Measured molecular weight of the two peaks was 157.9 kDa (theoretical hexamer = 160.0 kDa) and 80.0 kDa (theoretical trimer = 80.0 kDa).

## Supplemental Note

Sequence of UC Berkeley Macrolab plasmid 2AT (Addgene #29665), used for NucC plasmid digestion assays and high-throughput sequencing:

TTCTTGAAGACGAAAGGGCCTCGTGATACGCCTATTTTTATAGGTTAATGTCATGATAATAATGGTTTCTTAGACGTCAGGTGGCACTTTTCGGGGAAATGTGCGCGGAACCCCTATTTGTTTATTTTTCTAAATACATTCAAATATGTATCCGCTCATGAGACAATAACCCTGATAAATGCTTCAATAACATTGAAAAAGGAAGAGTATGAGTATTCAACATTTCCGTGTCGCCCTTATTCCCTTTTTTGCGGCATTTTGCCTTCCTGTTTTTGCTCACCCAGAAACGCTGGTGAAAGTAAAAGATGCTGAAGATCAGTTGGGTGCACGAGTGGGTTACATCGAACTGGATCTCAACAGCGGTAAGATCCTTGAGAGTTTTCGCCCCGAAGAACGTTTTCCAATGATGAGCACTTTTAAAGTTCTGCTATGTGGCGCGGTATTATCCCGTGTTGACGCCGGGCAAGAGCAACTCGGTCGCCGCATACACTATTCTCAGAATGACTTGGTTGAGTACTCACCAGTCACAGAAAAGCATCTTACGGATGGCATGACAGTAAGAGAATTATGCAGTGCTGCCATAACCATGAGTGATAACACTGCGGCCAACTTACTTCTGACAACGATCGGAGGACCGAAGGAGCTAACCGCTTTTTTGCACAACATGGGGGATCATGTAACTCGCCTTGATCGTTGGGAACCGGAGCTGAATGAAGCCATACCAAACGACGAGCGTGACACCACGATGCCTGCAGCAATGGCAACAACGTTGCGCAAACTATTAACTGGCGAACTACTTACTCTAGCTTCCCGGCAACAATTAATAGACTGGATGGAGGCGGATAAAGTTGCAGGACCACTTCTGCGCTCGGCCCTTCCGGCTGGCTGGTTTATTGCTGATAAATCTGGAGCCGGTGAGCGTGGGTCTCGCGGTATCATTGCAGCACTGGGGCCAGATGGTAAGCCCTCCCGTATCGTAGTTATCTACACGACGGGGAGTCAGGCAACTATGGATGAACGAAATAGACAGATCGCTGAGATAGGTGCCTCACTGATTAAGCATTGGTAACTGTCAGACCAAGTTTACTCATATATACTTTAGATTGATTTAAAACTTCATTTTTAATTTAAAAGGATCTAGGTGAAGATCCTTTTTGATAATCTCATGACCAAAATCCCTTAACGTGAGTTTTCGTTCCACTGAGCGTCAGACCCCGTAGAAAAGATCAAAGGATCTTCTTGAGATCCTTTTTTTCTGCGCGTAATCTGCTGCTTGCAAACAAAAAAACCACCGCTACCAGCGGTGGTTTGTTTGCCGGATCAAGAGCTACCAACTCTTTTTCCGAAGGTAACTGGCTTCAGCAGAGCGCAGATACCAAATACTGTCCTTCTAGTGTAGCCGTAGTTAGGCCACCACTTCAAGAACTCTGTAGCACCGCCTACATACCTCGCTCTGCTAATCCTGTTACCAGTGGCTGCTGCCAGTGGCGATAAGTCGTGTCTTACCGGGTTGGACTCAAGACGATAGTTACCGGATAAGGCGCAGCGGTCGGGCTGAACGGGGGGTTCGTGCACACAGCCCAGCTTGGAGCGAACGACCTACACCGAACTGAGATACCTACAGCGTGAGCTATGAGAAAGCGCCACGCTTCCCGAAGGGAGAAAGGCGGACAGGTATCCGGTAAGCGGCAGGGTCGGAACAGGAGAGCGCACGAGGGAGCTTCCAGGGGGAAACGCCTGGTATCTTTATAGTCCTGTCGGGTTTCGCCACCTCTGACTTGAGCGTCGATTTTTGTGATGCTCGTCAGGGGGGCGGAGCCTATGGAAAAACGCCAGCAACGCGGCCTTTTTACGGTTCCTGGCCTTTTGCTGGCCTTTTGCTCACATGTTCTTTCCTGCGTTATCCCCTGATTCTGTGGATAACCGTATTACCGCCTTTGAGTGAGCTGATACCGCTCGCCGCAGCCGAACGACCGAGCGCAGCGAGTCAGTGAGCGAGGAAGCGGAAGAGCGCCTGATGCGGTATTTTCTCCTTACGCATCTGTGCGGTATTTCACACCGCATATATGGTGCACTCTCAGTACAATCTGCTCTGATGCCGCATAGTTAAGCCAGTATACACTCCGCTATCGCTACGTGACTGGGTCATGGCTGCGCCCCGACACCCGCCAACACCCGCTGACGCGCCCTGACGGGCTTGTCTGCTCCCGGCATCCGCTTACAGACAAGCTGTGACCGTCTCCGGGAGCTGCATGTGTCAGAGGTTTTCACCGTCATCACCGAAACGCGCGAGGCAGCTGCGGTAAAGCTCATCAGCGTGGTCGTGAAGCGATTCACAGATGTCTGCCTGTTCATCCGCGTCCAGCTCGTTGAGTTTCTCCAGAAGCGTTAATGTCTGGCTTCTGATAAAGCGGGCCATGTTAAGGGCGGTTTTTTCCTGTTTGGTCACTGATGCCTCCGTGTAAGGGGGATTTCTGTTCATGGGGGTAATGATACCGATGAAACGAGAGAGGATGCTCACGATACGGGTTACTGATGATGAACATGCCCGGTTACTGGAACGTTGTGAGGGTAAACAACTGGCGGTATGGATGCGGCGGGACCAGAGAAAAATCACTCAGGGTCAATGCCAGCGCTTCGTTAATACAGATGTAGGTGTTCCACAGGGTAGCCAGCAGCATCCTGCGATGCAGATCCGGAACATAATGGTGCAGGGCGCTGACTTCCGCGTTTCCAGACTTTACGAAACACGGAAACCGAAGACCATTCATGTTGTTGCTCAGGTCGCAGACGTTTTGCAGCAGCAGTCGCTTCACGTTCGCTCGCGTATCGGTGATTCATTCTGCTAACCAGTAAGGCAACCCCGCCAGCCTAGCCGGGTCCTCAACGACAGGAGCACGATCATGCGCACCCGTGGCCAGGACCCAACGCTGCCCGAGATGCGCCGCGTGCGGCTGCTGGAGATGGCGGACGCGATGGATATGTTCTGCCAAGGGTTGGTTTGCGCATTCACAGTTCTCCGCAAGAATTGATTGGCTCCAATTCTTGGAGTGGTGAATCCGTTAGCGAGGTGCCGCCGGCTTCCATTCAGGTCGAGGTGGCCCGGCTCCATGCACCGCGACGCAACGCGGGGAGGCAGACAAGGTATAGGGCGGCGCCTACAATCCATGCCAACCCGTTCCATGTGCTCGCCGAGGCGGCATAAATCGCCGTGACGATCAGCGGTCCAGTGATCGAAGTTAGGCTGGTAAGAGCCGCGAGCGATCCTTGAAGCTGTCCCTGATGGTCGTCATCTACCTGCCTGGACAGCATGGCCTGCAACGCGGGCATCCCGATGCCGCCGGAAGCGAGAAGAATCATAATGGGGAAGGCCATCCAGCCTCGCGTCGCGAACGCCAGCAAGACGTAGCCCAGCGCGTCGGCCGCCATGCCGGCGATAATGGCCTGCTTCTCGCCGAAACGTTTGGTGGCGGGACCAGTGACGAAGGCTTGAGCGAGGGCGTGCAAGATTCCGAATACCGCAAGCGACAGGCCGATCATCGTCGCGCTCCAGCGAAAGCGGTCCTCGCCGAAAATGACCCAGAGCGCTGCCGGCACCTGTCCTACGAGTTGCATGATAAAGAAGACAGTCATAAGTGCGGCGACGATAGTCATGCCCCGCGCCCACCGGAAGGAGCTGACTGGGTTGAAGGCTCTCAAGGGCATCGGTCGACGCTCTCCCTTATGCGACTCCTGCATTAGGAAGCAGCCCAGTAGTAGGTTGAGGCCGTTGAGCACCGCCGCCGCAAGGAATGGTGCATGCAAGGAGATGGCGCCCAACAGTCCCCCGGCCACGGGGCCTGCCACCATACCCACGCCGAAACAAGCGCTCATGAGCCCGAAGTGGCGAGCCCGATCTTCCCCATCGGTGATGTCGGCGATATAGGCGCCAGCAACCGCACCTGTGGCGCCGGTGATGCCGGCCACGATGCGTCCGGCGTAGAGGATCGAGATCTCGATCCCGCGAAATTAATACGACTCACTATAGGGAGACCACAACGGTTTCCCTCTAGTGCCGGCTCCGGAGAGCTCTTTAATTAAGCGGCCGCCCTGCAGGACTCGAGTTCTAGAAATAATTTTGTTTAACTTTAAGAAGGAGATATAGATATCCCAACTCCATAAGGATCCGCGATCGCGGCGCGCCACCTGGTGGCCGGCCGGTACCACGCGTGCGCGCTGATCCGGCTGCTAACAAAGCCCGAAAGGAAGCTGAGTTGGCTGCTGCCACCGCTGAGCAATAACTAGCATAACCCCTTGGGGCCTCTAAACGGGTCTTGAGGGGTTTTTTGCTGAAAGGAGGAACTATATCCGGACATCCACAGGACGGGTGTGGTCGCCATGATCGCGTAGTCGATAGTGGCTCCAAGTAGCGAAGCGAGCAGGACTGGGCGGCGGCCAAAGCGGTCGGACAGTGCTCCGAGAACGGGTGCGCATAGAAATTGCATCAACGCATATAGCGCTAGCAGCACGCCATAGTGACTGGCGATGCTGTCGGAATGGACGACATCCCGCAAGAGGCCCGGCAGTACCGGCATAACCAAGCCTATGCCTACAGCATCCAGGGTGACGGTGCCGAGGATGACGATGAGCGCATTGTTAGATTTCATACACGGTGCCTGACTGCGTTAGCAATTTAACTGTGATAAACTACCGCATTAAAGCTTATCGATGATAAGCTGTCAAACATGAGAA

---

**Table S1.**
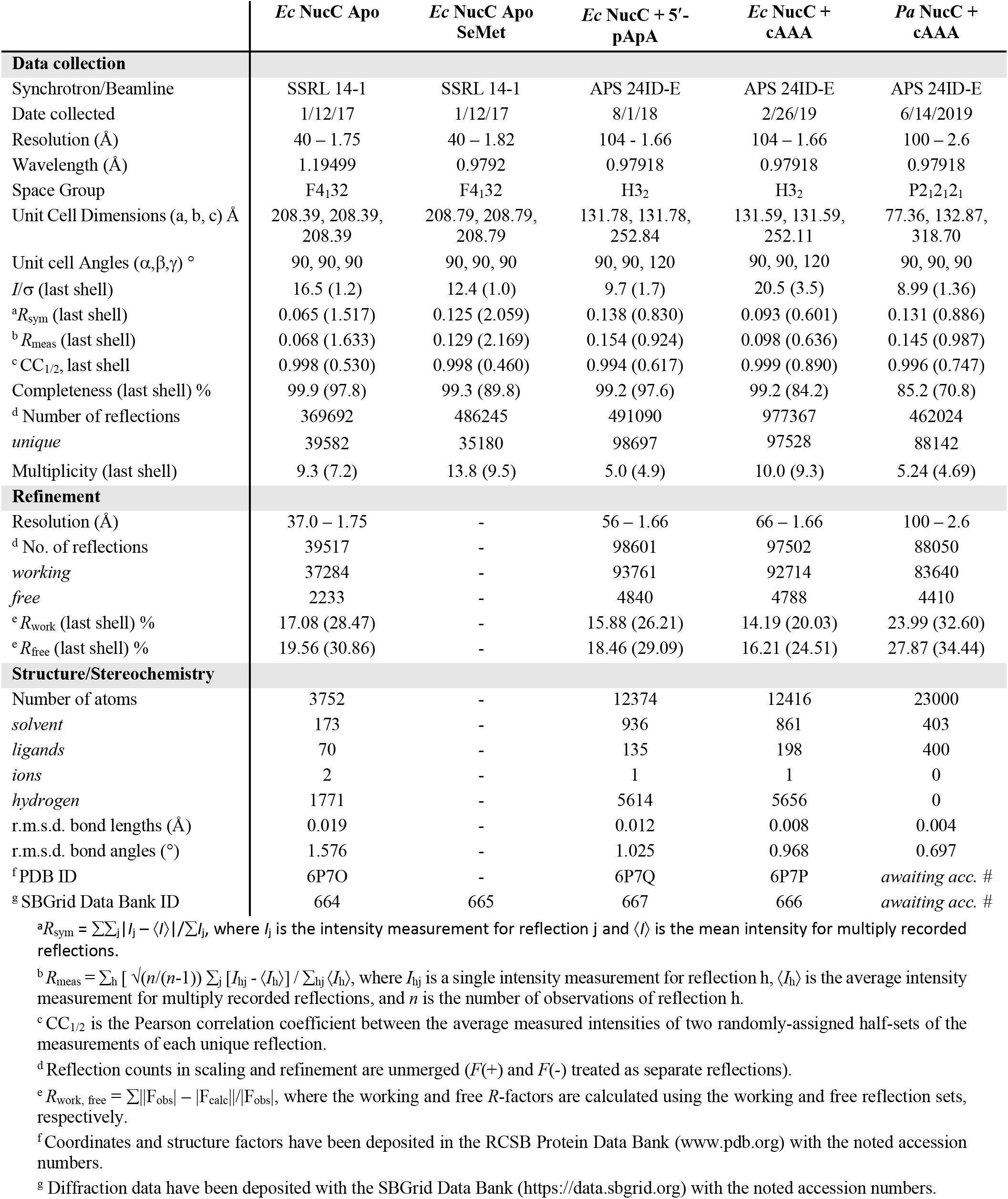
Crystallographic data collection and refinement

**Table S2.** NucC binding second messenger molecules in high-salt conditions

| Second messenger | $K_d$ (μM) | $N$ per NucC trimer<sup>a</sup> |
| --- | --- | --- |
| 3',3' cyclic di-AMP | 14.4 | 0.60 |
| 5'-pApA | 15.0 | 0.72 |
| 2',3' cyclic di-AMP | *no binding* | - |
| 3',3' cyclic di-GMP | *no binding* | - |
| 5'-pGpG | *no binding* | - |
| 3',3' cyclic GMP-AMP (cGAMP) | 9.3 | 0.75 |
| AMP | *no binding* | - |

<sup>a</sup> $N$ refers to the stoichiometry of second messenger binding to the NucC homotrimer.

**Table S3.**
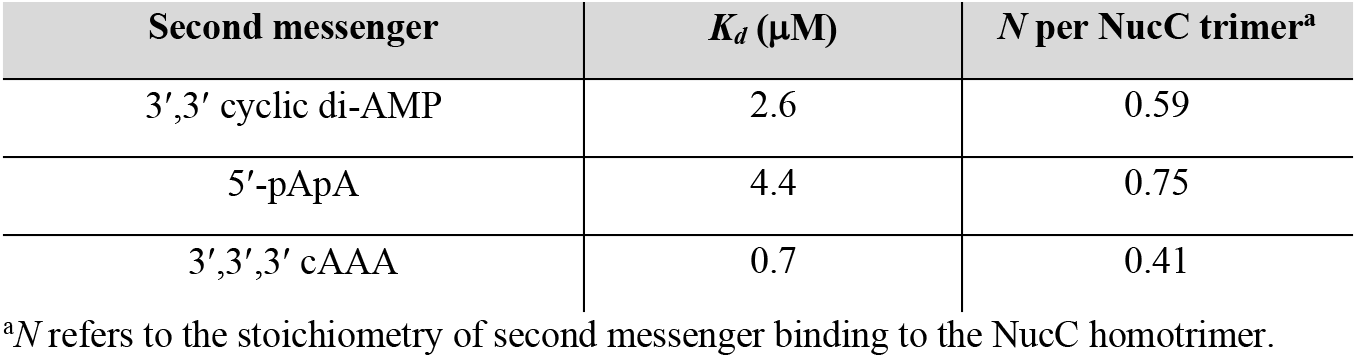
NucC binding second messenger molecules in low-salt conditions

| Second messenger | $K_d$ (μM) | $N$ per NucC trimer<sup>a</sup> |
| --- | --- | --- |
| 3',3' cyclic di-AMP | 2.6 | 0.59 |
| 5'-pApA | 4.4 | 0.75 |
| 3',3',3' cAAA | 0.7 | 0.41 |

<sup>a</sup> $N$ refers to the stoichiometry of second messenger binding to the NucC homotrimer.

**Table S4.**
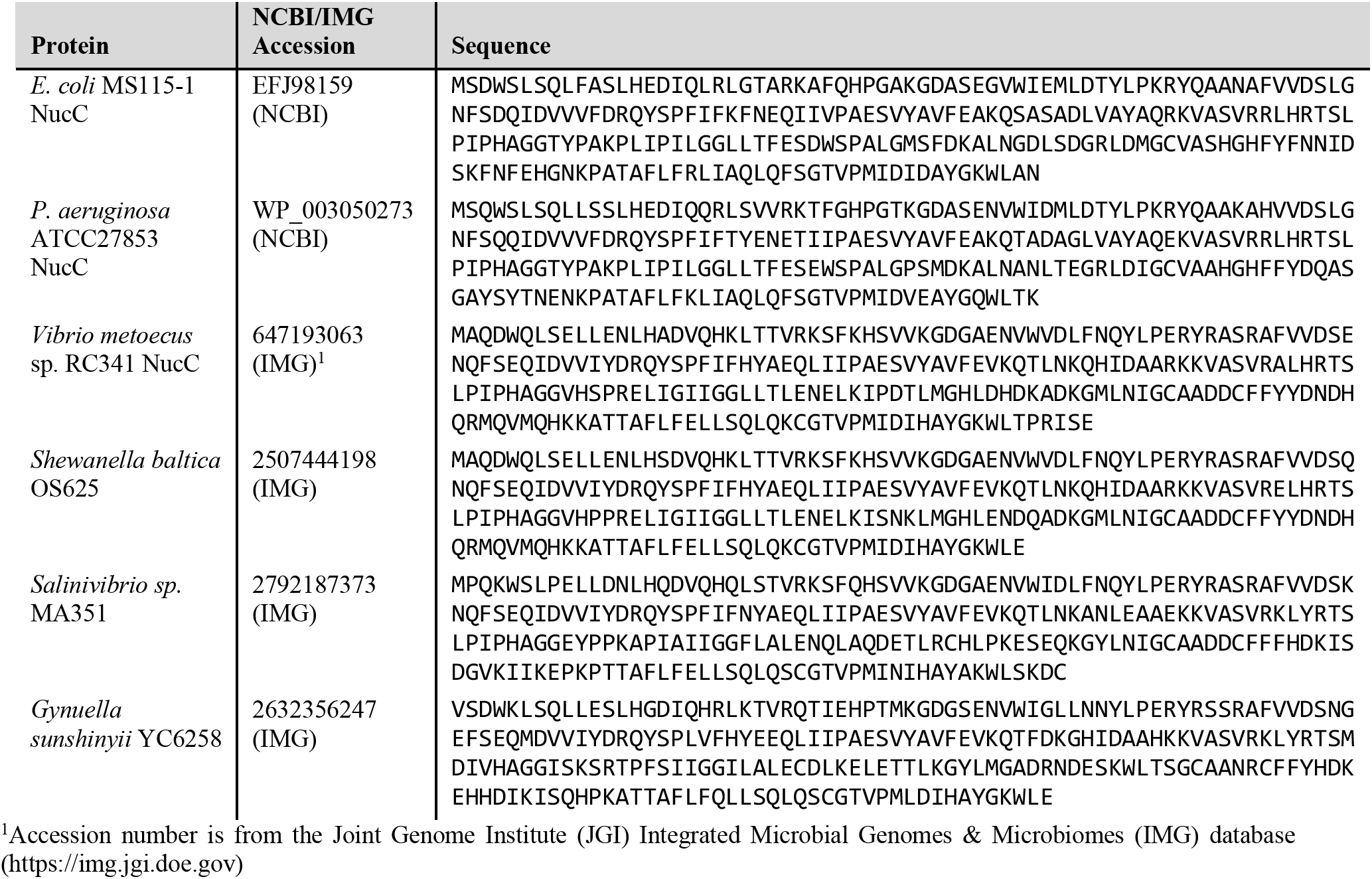
Protein Sequences

